# A Necrotizing Toxin Promotes *Pseudomonas syringae* Infection Across Evolutionarily Divergent Plant Lineages

**DOI:** 10.1101/2024.07.17.603760

**Authors:** Kristina Grenz, Khong-Sam Chia, Emma K. Turley, Alexa S. Tyszka, Rebecca E. Atkinson, Jacob Reeves, Martin Vickers, Martin Rejzek, Joseph F. Walker, Philip Carella

**Affiliations:** Cell and Developmental Biology, John Innes Centre, Norwich NR4 7UH, UK; Department of Biological Sciences, University of Illinois at Chicago, Chicago, IL 60607, USA; Computational and Systems Biology, John Innes Centre, Norwich NR4 7UH, UK; Molecular Microbiology, John Innes Centre, Norwich NR4 7UH, UK

**Author notes:** **Corresponding author:** Philip Carella. These authors contributed equally (Kristina Grenz, Khong-Sam Chia).

## Abstract

The *Pseudomonas syringae* species complex harbors a diverse range of plant pathogenic bacteria. While much of the current understanding of *P. syringae* is centered on interactions with flowering plants, much less is known about infection in evolutionarily divergent non-flowering lineages. Here, we took a comparative evolutionary approach to understand how *P. syringae* infects distantly related plants. We identify broad host *P. syringae* isolates causing significant disease in the liverwort *Marchantia polymorpha*, the fern *Ceratopteris richardii*, and the flowering plant *Nicotiana benthamiana*, which last shared a common ancestor over 500 million years ago. We demonstrate that phytotoxin enriched isolates belonging to the phylogroup 2 clade of the *P. syringae* species complex are particularly virulent in non-flowering plants, relying on a combination of type-3 secreted effector proteins and the lipopeptide phytotoxin syringomycin. The application of purified syringomycin promotes necrosis in diverse host tissues and activates conserved genes associated with redox regulation and cell death. Toxin-deficient phylogroups normally unable to infect *Marchantia* thalli exhibit enhanced bacterial growth when supplemented with exogenous syringomycin, further highlighting its role as a host-range defining factor in *Pseudomonas*. Collectively our research reveals a key role for the lipopeptide syringomycin in promoting *Pseudomonas* colonization, which works in concert with type-3 effector proteins to antagonize an exceptionally wide spectrum of land plants.

## INTRODUCTION

*Pseudomonas syringae* is a globally distributed gram-negative bacteria that causes disease on a wide array of flowering plants^1^. For decades it has been exploited to dissect key processes that underpin pathogen virulence and plant adaptation to infection in genetic model systems like *Arabidopsis thaliana* or in economically-relevant hosts like tomato or kiwi^2^. Evolving from a *P. fluorescens-*like predecessor over 150 million years ago, *P. syringae* has expanded into a monophyletic species complex with 13 distinct phylogroups (PGs) harboring 64 known pathovars and plant-pathogenic species like *P. viridiflava* and *P. cichorii* among others^3,4^. In general, the complex is divided into two groups, the late-branching clade of canonical *P. syringae* pathovars (PGs 1-6 and 10) that are historically associated with crop infection, and a smaller ‘secondary’ group of early-branching pathovars and species (PGs 7-9 and 11-13) typically isolated from abiotic environmental settings in addition to plants^4,5^. While much of our knowledge is derived from interactions with flowering plants, emerging ecological data suggests that *P. syringae* naturally associates with non-flowering plants^6^. However, the molecular processes enabling *P. syringae* infection across the full spectrum of plant evolution remain to be clarified.

*P. syringae* relies on a variety of virulence factors to promote disease, including effector proteins translocated into plant cells through the ‘needle-like’ type-3 secretion system (T3SS) and small-molecule phytotoxins. The *P. syringae* species complex encodes a diverse set of type-3 effector proteins displaying a range of enzymatic activities and immune suppressing functions^7^. Among this diversity, four effectors are present across the vast majority of phylogroups, including those within the conserved effector locus (CEL; AvrE, hopM1, and hopAA1) that are essential for plant infection^4,7,8^. By contrast, phytotoxins are sparsely distributed across the species complex, with exception to PG2 strains that deploy diverse toxins with roles in proteasome inhibition (syringolin)^9^, mimicking host hormones (coronatine, auxin)^10,11^, disrupting host metabolism (tabtoxin, mangotoxin, phaseolotoxin)^12–14^, or perforating host membranes (syringomycin and syringopeptin)^15^. Intriguingly, toxin-enriched *P. syringae* isolates within PG2 typically encode smaller type-3 effector repertoires, suggesting that the general action of phytotoxins may relieve the evolutionary pressure to accumulate type-3 effectors^4^.

Since their emergence onto land over 500 million years ago, plants have evolved into major lineages ranging from non-vascular/non-seed bryophytes (liverworts, mosses, and hornworts), vascular non-seed lycophytes and monilophytes (clubmosses and ferns), seed-bearing but non-flowering gymnosperms, and flowering seed plants^16^. Our current understanding of *P. syringae*-plant interactions is dominated by flowering plants, though natural associations between *P. syringae* complex members and non-flowering plants have been observed in mosses, ferns, and gymnosperms^6,17^. For example, the *P. syringae* pathovar (pv.) *cunninghamiae* naturally infects needles of the conifer (gymnosperm) *Cunninghamia lanceolata*^17^. In addition, commonly studied laboratory strains like *P. syringae pv. tomato* DC3000 can infect the model liverwort (bryophyte) *Marchantia polymorpha*, where it relies on the T3SS to support bacterial proliferation within the vegetative plant body (thallus)^18,19^. Collectively, these studies hint towards the idea that the *P. syringae* complex carries the potential to infect diverse and distantly related hosts across the full spectrum of plant evolution.

In this study, we interrogate diverse isolates from across the *P. syringae* species complex for their capacity to infect distantly related host plants *M. polymorpha* (liverwort), *Ceratopteris richardii* (fern), and *Nicotiana benthamiana* (flowering plant). We demonstrate that while individual isolates display differential disease potentials in different hosts, those from the PG2 clade exhibit the broadest experimental host range and are capable of infecting three divergent plant lineages that last shared a common ancestor over 450 million years ago. Comparative genomics and functional dissection of prevalent virulence factors in the representative PG2 isolate *P. syringae* pv. *syringae* B728a demonstrated key roles in virulence for both the T3SS and the lipopeptide toxin syringomycin, which itself was particularly important for non-flowering plant infection. Experiments leveraging purified syringomycin further highlighted its role in promoting host necrosis, which enhanced the *in planta* bacterial proliferation of toxin deficient phylogroups during liverwort infections. Collectively, our results highlight the broad role of the phytotoxin syringomycin and the T3SS in colonizing evolutionarily divergent host plants and solidifies *P. syringae* as an effective pathogen of the plant kingdom.

## RESULTS

### The *Pseudomonas syringae* species complex infects diverse land plant lineages

To determine the extent to which *P. syringae* infects evolutionarily divergent plants, we interrogated 53 diverse isolates for their ability to cause disease in the liverwort *M. polymorpha* (non-vascular, non-seed), the fern *C. richardii* (vascular, non-seed), and the angiosperm *N. benthamiana* (vascular, seed) (Fig. 1A). Where possible, a minimum of three representative ‘pathotype’ strains were tested for each major *P. syringae* phylogroup, in addition to *Pseudomonas species* that are either unassigned (unsequenced) or outside of the species complex but are potentially pathogenic in non-flowering plants like *P. asplenii* (originally sourced from diseased ferns)^20^ (Data S1). By quantifying bacterial proliferation within vegetative photosynthetic tissues (liverwort thallus, fern frond, angiosperm leaf) and monitoring disease symptoms, we determined that many isolates display differential virulence phenotypes across divergent hosts (Fig. 1B; Data S1, S2). For example, *Marchantia* liverworts are highly susceptible to *P. viridiflava, P. cichorii,* and *P. asplenii,* while the fern *Ceratopteris* succumbs only to *P. asplenii* and *Nicotiana* leaves resist all three species. By contrast, we observed that several PG2 isolates harbor an exceptionally broad disease potential, such that high bacterial growth levels and strong disease symptoms were observed in all three evolutionarily divergent hosts. Collectively, these data demonstrate that *P. syringae* and related *Pseudomonads* are indeed efficient pathogens of non-flowering and flowering plants alike.

**Figure 1.**
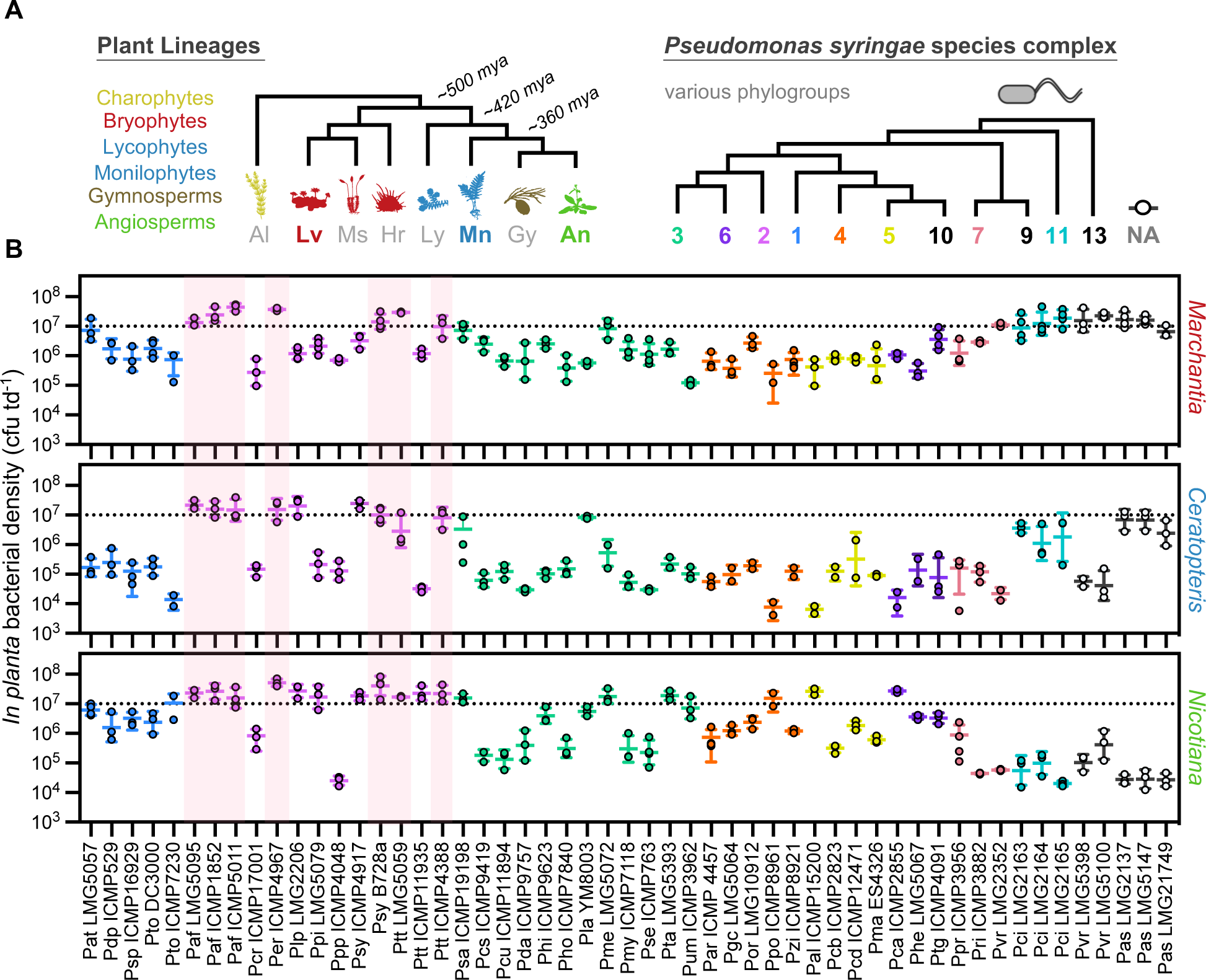
*Pseudomonas syringae* infects evolutionarily divergent plant lineages. (A) Schematic representation of diverse plant lineages and the *Pseudomonas syringae* (*Psy*) species complex. Plants from diverse lineages are depicted with cartoons representing algae (Al), bryophytes (Lv, liverworts; Ms, mosses; Hr, hornworts), lycophytes (Ly), monilophytes (Mn), gymnosperms (Gy), and angiosperms (An) at the approximate timescales (mya, millions of years ago). Major phylogroups (PGs) of the *Psy* species complex are indicated (NA, not associated/determined). (B) Bacterial proliferation in the liverwort *Marchantia polymorpha,* fern *Ceratopteris richardii,* and angiosperm *Nicotiana benthamiana. In planta* bacterial densities are expressed as colony forming units (CFU) per tissue disc (td). Individual datapoints represent mean bacterial densities for a given experimental replicate, with at least two experimental replicates performed per *Psy* isolate. Bacterial densities were determined 3 days post infection (dpi) for *Marchantia* and *Ceratopteris,* or 2 dpi for *Nicotiana.* A horizontal line represents the mean of all experimental replicates, with error bars indicating standard deviation. PG2 isolates infecting all three hosts are highlighted in pink.

### *P. syringae* virulence factors are similarly upregulated within divergent hosts

To better understand how *P. syringae* manipulates diverse plant hosts, we interrogated bacterial genomes for the presence of major virulence determinants like type-3 secreted effector proteins and phytotoxins (Fig. 2A). By leveraging existing genomic resources, we assessed virulence factor prevalence in sets of infectious isolates (on *Marchantia, Ceratopteris, Nicotiana,* or “All hosts”) relative to the total set of tested isolates (Fig. 2; Fig. S1A). In general, phytotoxins were enriched in isolates infecting each non-flowering plant or in all three hosts, with the exception of coronatine and tabtoxin (Fig. 2B; Data S1). While conserved type-3 effector families like AvrE, hopM1, and hopAA1 were present in all tested isolates, accessory effectors displayed differential abundances across our comparisons (Fig. 2C; Fig. S1B; Data S1). Notably, hopQ1 was less prevalent in isolates infecting *Nicotiana* and other plants relative to the total set of tested isolates, which is consistent with its known function as an avirulence effector in *N. benthamiana*^21^.

**Figure 2.**
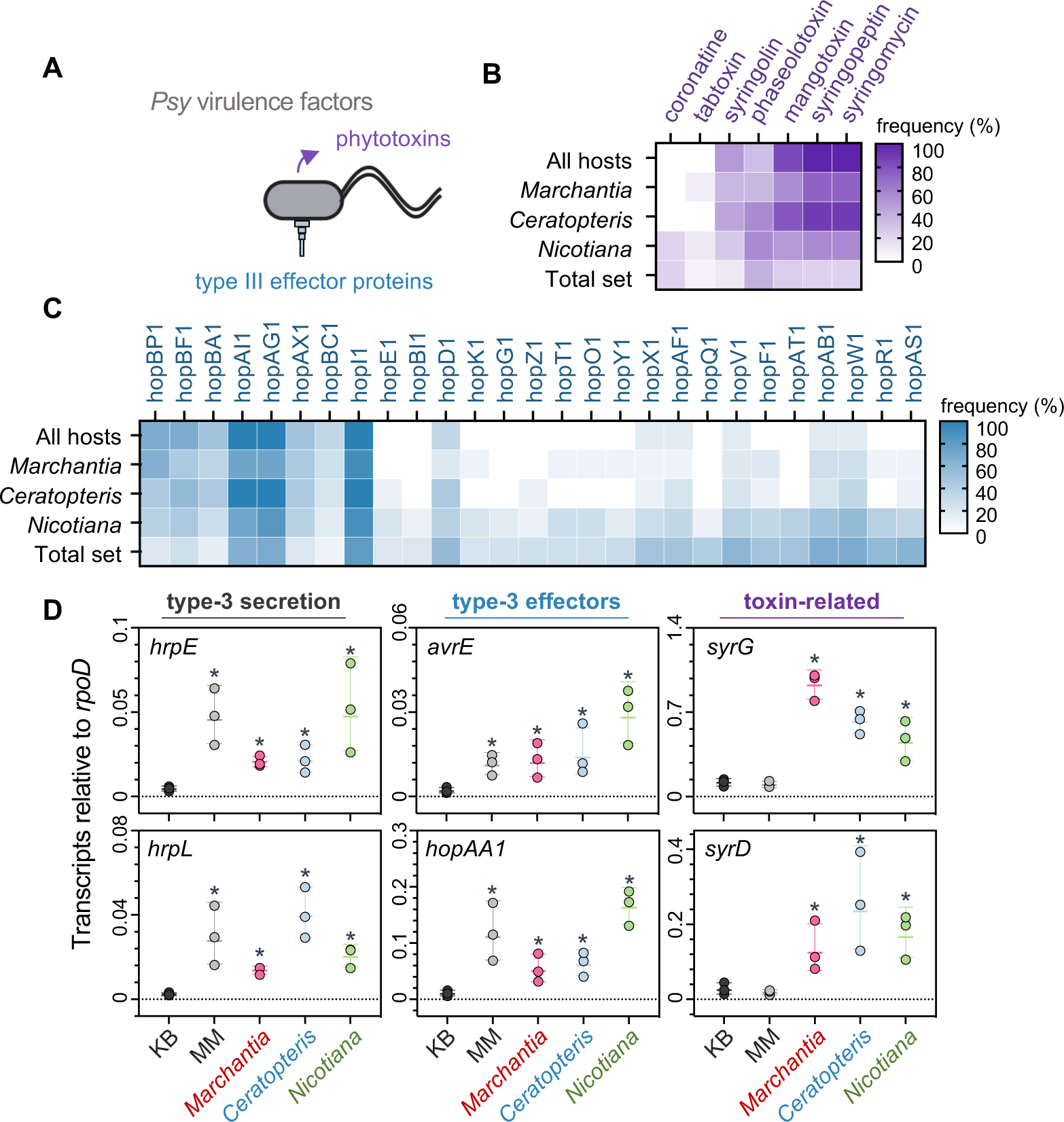
*Pseudomonas* virulence factors are commonly deployed *in planta*. (A) Schematic demonstrating the two main classes of *Pseudomonas syringae* virulence factors; type-3 effector proteins secreted through the type-3 secretion system (T3SS), and phytotoxins. (B) Prevalence of major phytotoxins in sets of *Pseudomonas syringae* isolates infecting all three hosts (all hosts), the liverwort *Marchantia polymorpha* (*Marchantia*), the fern *Ceratopteris richardii* (*Ceratopteris*), the flowering plants *Nicotiana benthamiana* (*Nicotiana*), or the total set of all tested isolates (total set). Prevalence is shown as the percentage of strains with a given toxin relative to the total number of isolates within a given set. (C) Prevalence of type-3 effector proteins in sets of *Pseudomonas syringae* isolates infecting all hosts, *Marchantia*, *Ceratopteris*, *Nicotiana*, or the total set. Prevalence is shown as the percentage of strains with a given toxin relative to the total number of isolates within a given set. Only effectors with at least 20% difference in abundance between any two sets are shown here. (D) qRT-PCR analysis of key *P. syringae* pv. *syringae* B728a virulence factors expressed in rich media (KB, King’s B), minimal media (MM), or *in planta* during diverse host infections. Expression levels of genes associated with the T3SS (*hrpE, hrpL*), conserved effectors (*avrE, hopAA1*), or the phytotoxin syringomycin (*syrD, syrG*) are shown relative to the internal *rpoD* control. Each datapoint represents pooled tissue (n ≥ 10) from three independent experiments. Expression was quantified in 1-day old liquid cultures or 6 hours post infection *in planta.* An asterisk (*) denotes statistically significant differences in transcript abundance in comparison to the KB control (Student’s t-test, p < 0.05). Error bars represent standard deviation.

Next, we performed quantitative reverse transcription PCR (qRT-PCR) to assess expression levels of *P. syringae* virulence genes during diverse host plant infections. To achieve this, we used the model pathovar *P. syringae* pv. *syringae* B728a, as it was one of seven PG2 isolates capable of infecting *Marchantia, Ceratopteris,* and *Nicotiana*. We focused on genes involved in the regulation or function of the T3SS (*hrpE*, *hrpL*), effectors within the CEL (*avrE*, *hopAA1*), and phytotoxin regulation or secretion (*syrG*, *syrD*). Expression levels were also analyzed during growth in rich media (KB) where virulence factor gene expression is not activated, and in minimal media (MM) that mimics the plant apoplast and activates expression^22^. Relative to the KB media control, bacteria grown in MM upregulated T3SS genes and effectors, whereas toxin-related genes failed to activate in our conditions (Fig. 2D). Importantly, T3SS components, effectors, and phytotoxin gene expression were all significantly upregulated *in planta* irrespective of host lineage, supporting the idea that *P. syringae* utilize a common virulence program in divergent host plants.

### A lipopeptide phytotoxin and type-3 effectors underpin B728a virulence in divergent hosts

Given that effectors and toxins are upregulated during bacterial infection of distantly related host plants, we next examined their contributions to disease establishment. Using *P. syringae* B728a as a model, we challenged *Marchantia, Ceratopteris,* and *Nicotiana* with wild-type (B728a^WT^) versus T3SS-defective ΔhrcC mutants (unable to translocate effectors)^23^ or with ΔsyrP mutants unable to produce the lipopeptide phytotoxin syringomycin^24^. In each lineage, wild-type B728a^WT^ exhibited prominent disease symptoms and reached high *in planta* bacterial densities, whereas ΔhrcC mutants failed to cause disease symptoms or appreciably proliferate *in planta* (Fig. 3AB). Intriguingly, the ΔsyrP mutant exhibited minimal disease symptoms in *N. benthamiana* leaves but grew to high densities similar to B728a^WT^, whereas non-flowering plants displayed minimal disease symptoms but supported low levels of *in planta* bacterial growth. These results indicate that while the phytotoxin syringomycin is essential for the full manifestation of disease, its role in supporting bacterial growth is more important in non-flowering *Marchantia* and *Ceratopteris* compared to the angiosperm *Nicotiana*.

**Figure 3.**
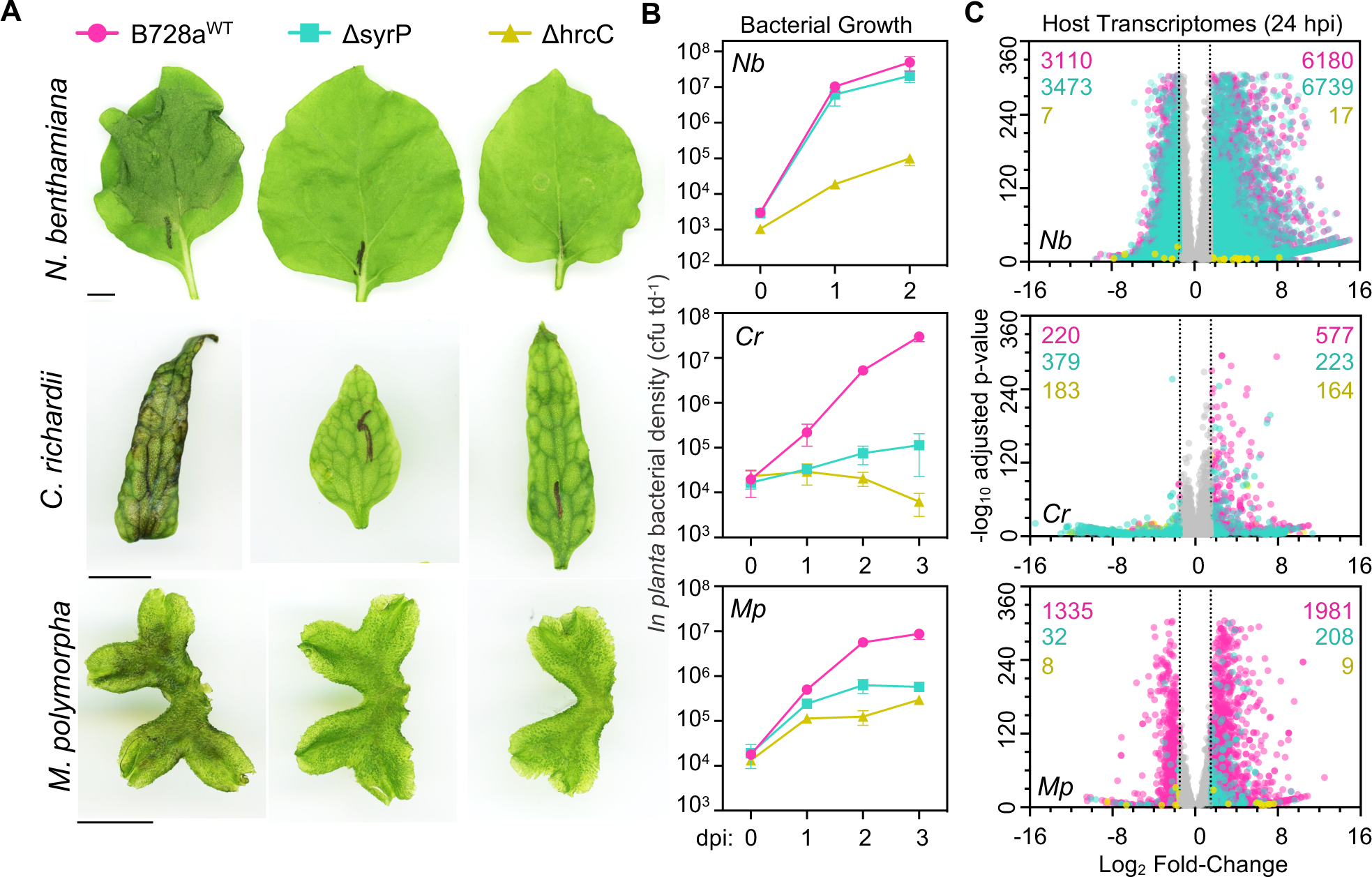
Effectors and syringomycin are essential for virulence in non-flowering plants. (A) Symptom development of diverse plants infected with *P. syringae* pv. *syringae* B728a. *Marchantia, Ceratopteris,* and *Nicotiana* were infected with wildtype (B728a^WT^), ΔsyrP, and ΔhrcC mutants. Symptoms were recorded 3 days post infection (dpi) for *Marchantia* and *Ceratopteris,* or 2 dpi for *Nicotiana* (n ≥ 8 for all plants and treatments). This experiment was performed at least three times with similar results. Scale bars = 1 cm. (B) *In planta* bacterial growth curves of B728a^WT^, ΔsyrP, and ΔhrcC in diverse host plants *Marchantia* (*Mp*), *Ceratopteris* (*Cr*), and *Nicotiana* (*Nb*). Bacterial densities (CFU per tissue disc) were measured at the indicated dpi. Each datapoint represents the mean of three biological replicates (n ≥ 8 plants per treatment/timepoint), with error bars representing standard deviation. This experiment was performed 3 times with similar results. (C) Host transcriptome analyses of diverse plants infected with *P. syringae* at 24 hours post infection (hpi). Volcano plots display pairwise differential expression analysis per *Psy* genotype versus a mock-treated control in a given host. Significantly differentially expressed genes are displayed, with total numbers of differentially expressed genes for B728a^WT^, ΔsyrP, and ΔhrcC shown in descending order.

To better understand the impact that syringomycin and effectors exert on land plants, we performed RNA-sequencing (RNA-seq) experiments comparing the infection-associated expression profiles elicited by wild-type B728a^WT^, ΔsyrP, or ΔhrcC relative to mock-treated controls at 24 hours post inoculation in each host. Differential gene expression analyses identified prominent shifts in the transcriptomes of B728a^WT^ infected hosts across non-flowering and flowering plants alike, while responses to ΔhrcC mutants were minimal (Fig. 3C; Fig. S2; Data S3). Infection with the syringomycin-deficient ΔsyrP mutant had a greater overall impact on the *Nicotiana* transcriptome relative to *Marchantia* or *Ceratopteris,* consistent with the fact that ΔsyrP bacteria grow to high levels only in *Nicotiana* leaves. We then compared transcriptomes amongst the three divergent plants to delineate common molecular responses to *P. syringae* infection. Using established sets of orthologous protein coding genes (orthogroups) identified by OrthoFinder^25,26^, we determined orthologous genes that were consistently differentially expressed during infection across *Nicotiana, Ceratopteris,* and *Marchantia*. This revealed 56 orthogroups shared amongst diverse hosts during B728a^WT^ infection, 6 orthogroups elicited by ΔsyrP, and no overlap in ΔhrcC-elicited genes (Fig. S2D). Functional enrichment analyses for orthogroups responsive to disease establishment by B728a^WT^ demonstrated the common activation of ABC transporters, phenylpropanoid-related enzymes, calcium binding proteins, redox regulators, and transcription factors, whereas downregulated loci included cell wall enzymes/proteins and redox regulators (Data S3). Though functional enrichment analyses were not possible for the 6 orthogroups commonly responsive to ΔsyrP infection, we similarly observed the differential expression of peroxidases, ABC transporters, phenylalanine ammonia lyases (PAL), chalcone synthases (CHS), and a putative beta-lactamase (hydrolase).

### Syringomycin provokes host cell death in diverse plant lineages

Cyclic lipopeptides like syringomycin often display toxic activities across diverse eukaryotes like plants, algae, oomycetes, and fungi^27^. To explore the impact of syringomycin in non-flowering plants, we first purified the toxin from the syringomycin E (SRE) overproducing strain *P. syringae* pv. *syringae* isolate (B310D)^28^. Using an activity guided approach leveraging the inhibitory effect of syringomycin against the oomycete *Phytophthora infestans,* we obtained pure (>94%) and bioactive SRE (Fig. S3). We then assessed the impact of ectopic SRE application in all three divergent host plants by applying increasingly concentrated doses of the toxin. Mock-treated (water) controls and lower doses of SRE (0.01-0.025 mg ml^−1^) generally exhibited little impact on plant tissues, whereas moderate-to-higher doses of SRE (0.1-0.5 mg ml^−1^) led to visible cell death with complete tissue collapse (Fig. 4A; Fig. S4). The fern *C. richardii* showed the highest sensitivity to SRE, with prominent tissue collapse observed at lower doses (0.05 mg ml^−1^) relative to other plants (Fig. S4B). Next, we assessed the molecular responses of plants to SRE application using qRT-PCR. We focused on conserved marker genes belonging to orthogroups commonly responsive to B728a^WT^ infection that are associated with redox (OG68; *LOX1, lipoxygenase*) and cell-death (OG126; *AAA, AAA^+^-type ATPase*) processes in *Marchantia.* Relative to mock-treated controls, all *LOX1* and *AAA* orthologs were strongly responsive to SRE application (Fig. 4B). Together, these data support the idea that SRE imparts a necrotic phytotoxic effect on non-flowering and flowering plants alike.

**Figure 4.**
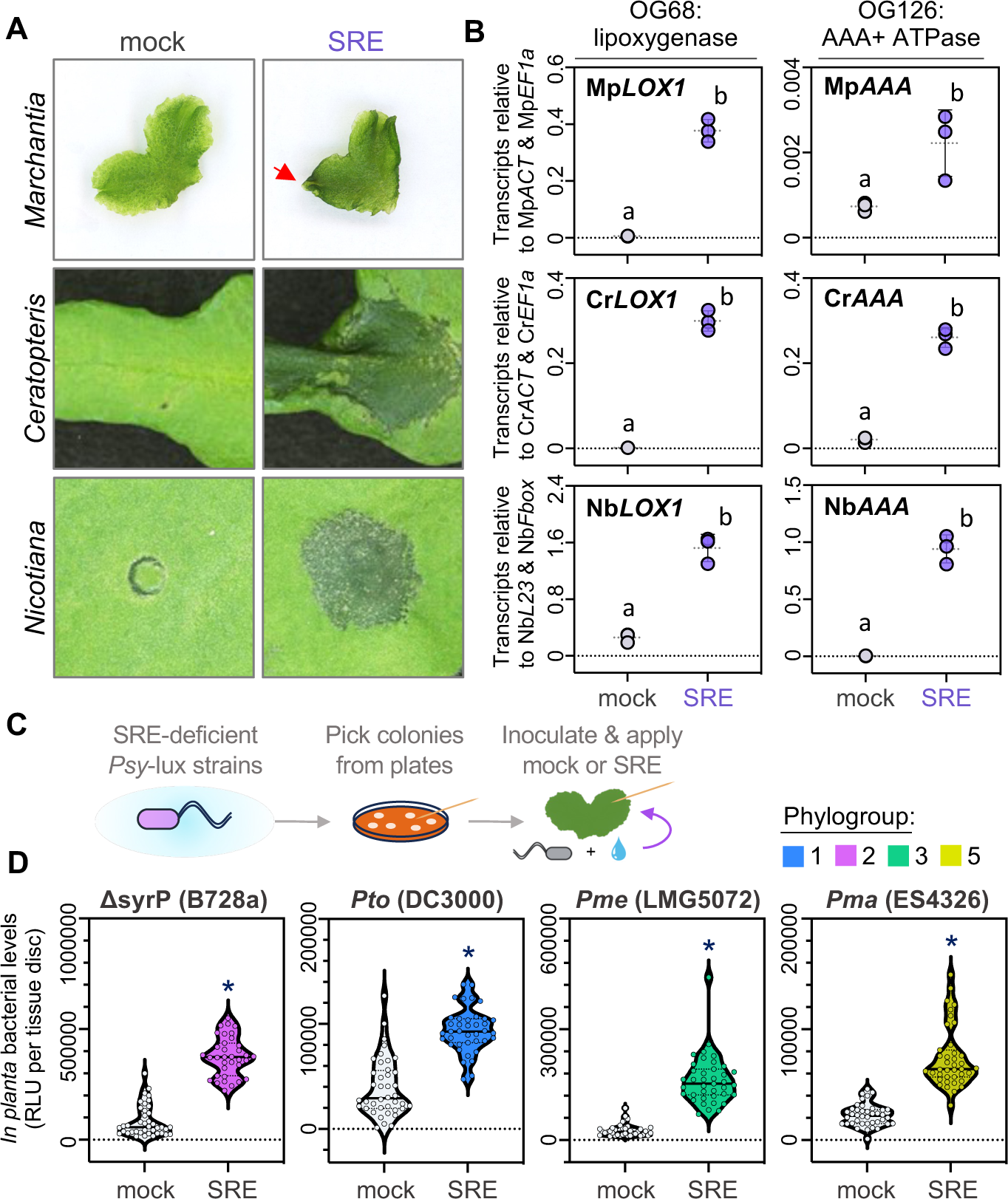
Syringomycin promotes necrotic cell death that extends *P. syringae* host range. (A) Macroscopic phenotypes of *Marchantia, Ceratopteris,* and *Nicotiana* treated with purified syringomycin E (SRE, 0.5 μg μl^−1^) compared to mock-treated (water) control. Images were taken 24 hours post treatment (n ≥ 5). A red arrow indicates a collapsed apical notch of *Marchantia.* This experiment was performed three times with similar results. (B) qRT-PCR analysis of orthogroups (OG) representing lipoxygenases (OG68: LOX1) or AAA+ATPases (OG126: AAA) in *Marchantia* (*Mp*)*, Ceratopteris* (*Cr*), and *Nicotiana* (*Nb*) at 24 hours post SRE application (0.5 μg μl^−1^) or mock-treatment (water). Expression values are shown relative to internal controls in each plant (Mp*ACT* & Mp*EF1a*; Cr*ACT* & Cr*EF1a*; Nb*L23* & Nb*Fbox*). Different letters signify statistically significant differences in transcript abundance (ANOVA, Tukey’s HSD, p < 0.05). Error bars represent standard deviation. Experiments were performed three times with similar results. (C) Experimental schematic outlining SRE rescue assays whereby bioluminescent reporter bacteria are grown on plates, colonies are picked using toothpicks, bacteria are wound-inoculated into *Marchantia* thalli, and then inoculation sites are provided with SRE or a mock treatment (water). (D) Luminescence quantification of SRE-deficient *P. syringae* isolates belonging to various phylogroups (PG). Data points represent relative luminescence units (RLU) per inoculation site (tissue disc) of 4-week-old *M. polymorpha* thalli wound-inoculated with bioluminescent ΔsyrP (PG2: *P. syringae* pv. *syringae* B728a), *Pto* (PG1: *P. syringae* pv. tomato DC3000), *Pme* (PG3: *P. syringae* pv. mellea LMG5072), or *Pma* (PG5: *P. syringae* pv. *maculicola* ES4326) and supplemented with water (mock) or syringomycin E (SRE, 0.5 μg μl^−1^). Violin plots include solid horizontal lines indicating median values (n ≥ 40 tissue discs per treatment, collected from at least 10 individual plants). An asterisk (*) denotes statistically significant differences in luminescence between SRE and mock-treated inoculations (Student’s t-test, p < 0.05). This experiment was performed three times with similar results.

### Ectopic syringomycin enhances the virulence of toxin-deficient *P. syringae* in *Marchantia*

Since syringomycin is a prominent *P. syringae* virulence factor associated with successful infection of non-flowering plants, we next addressed whether ectopic SRE improves the fitness of toxin-deficient *P. syringae* isolates. To accomplish this, we designed an infection scheme where *Marchantia* thalli were wound-inoculated with bioluminescent *P. syringae* isolates, ectopically supplied with purified SRE or a mock (water) control, and bacterial levels at the infection site were determined after 2 days (Fig. 4C). First, we established that wound-inoculation of the SRE-producing PG2 isolate *P. syringae* pv. *syringae* B728a^WT^ produced strong disease that correlated with bioluminescence, whereas the ΔsyrP B728a mutant and SRE non-producing isolates (PG1, Pto DC3000; PG3, Pme LMG5072; PG5, Pma ES4326) all failed to infect liverwort thalli (Fig. S4A). After confirming that wounding followed by SRE or water application did not produce luminescence (Fig. S4B), we quantified bacterial luminescence in toxin vs mock-treated inoculation sites. Across all tested isolates, ectopic SRE significantly increased bacterial densities relative to mock-treated controls (Fig. 4D; Fig. S4C). While the data demonstrate that SRE rescues the syringomycin-deficient ΔsyrP B728a mutant, it reveals that SRE promotes the *in planta* proliferation of diverse pathovars from across the *P. syringae* species during infection in a non-flowering plant.

## DISCUSSION

Our study highlights the broad disease potential harbored within the *P. syringae* species complex. By leveraging the divergent evolutionary histories of the liverwort *M. polymorpha,* the fern *C. richardii,* and the angiosperm *N. benthamiana,* we demonstrate the robust nature of *P. syringae* virulence, which manifests across the spectrum of plant evolution. Given that these hosts last shared a common ancestor over 500 million years ago^16^, we hypothesize that *P. syringae* virulence is centered on fundamental plant processes shared amongst the plant kingdom. By contrast, host responses to *P. syringae* likely diverged, as we observed limited overlap between infection transcriptomes of *Marchantia, Ceratopteris,* and *Nicotiana.* Despite this, we identified a core set of orthologous differentially expressed genes involved in phenylpropanoid metabolism, membrane transport, redox regulation, and transcriptional regulation. Among these processes, the upregulation of the MpMyb14 transcription factor alongside conserved phenylpropanoid metabolism genes like PAL and CHS is consistent with previous efforts to dissect biochemical defenses in *Marchantia*^29^, and suggests overlap between liverwort responses to oomycete and bacterial pathogens. The data also hint towards the ancestral roles of defense-related transcription factors (NAC and WRKY), whose immune functions remain to be dissected in non-flowering plants but are well established in angiosperms^30^. Collectively, our work reveals conserved transcriptomic signatures indicative of an ancestral program mediating host-microbe interactions.

Members of the *P. syringae* PG2 clade are the most broadly infectious across all three hosts. In particular, toxin-enriched strains exhibited the strongest disease phenotypes in both flowering and non-flowering plants, whereas toxin-deficient strains were generally unable to infect non-flowering plants. These results are consistent with previous research demonstrating the broad host virulence of PG2 isolates in angiosperms, adding functional context to emerging ecological studies demonstrating natural associations with non-flowering plants^6,31^. Isolates outside of PG2 demonstrated differential virulence profiles in divergent hosts, indicative of a continuum of compatibilities that are typical across plant-*Pseudomonas* interactions^4,31^. Whether these interactions are defined by the presence/absence of important virulence or avirulence factors remains to be determined.

Type-3 effector proteins play an essential role in *P. syringae* virulence across diverse hosts^7^. This is supported by our data and others showing that T3SS-deficient mutants unable to translocate effectors are not infectious in land plants^18,19^. Amongst the diversity of *P. syringae* effectors, the widely distributed CEL effectors AvrE and hopM1 target conserved host proteins to establish an aqueous environment essential for bacterial survival in the plant apoplast^32–34^. Recent structural, biochemical, and cell physiological investigation of AvrE activity in *Xenopus* oocyetes and plants also supports a direct role in mediating water and solute transfer across eukaryotic membranes^35^. Considering their fundamental requirement, it is likely that CEL effectors manipulate deeply conserved host proteins/processes to promote infection. While this remains to be verified for AvrE and hopM1, the conserved targeting of plant BAK1 orthologs by the *Pto* DC3000 effectors AvrPto and AvrPtoB supports the idea that *P. syringae* hijacks core plant processes^18,36^.

The cyclic lipopeptide toxin syringomycin is an amphiphilic molecule that forms ion-permeable pores in cell membranes to disrupt ion homeostasis and cause host tissue necrosis^15,28,37^. In flowering plants, syringomycin and the related toxin syringopeptin are broadly associated with pathogenesis, both within and outside of the *P. syringae* species complex. For example, both lipopeptide toxins underpin *P. fluorescens* transitions between pathogenic and commensal lifestyles on *Arabidopsis* roots, such that toxin-deficient mutants phenocopy commensal strains^38^. Moreover, *P. syringae* (PG2) interaction studies in *Prunus avium* (cherry) demonstrate a key role for syringomycin in promoting disease in fruit^28,39^, however toxins appear less important for infection in leaves or woody tissues where type-3 effectors play a more dominant role^39^. We similarly observed a limited role for syringomycin during angiosperm (*N. benthamiana*) leaf infections, yet the toxin was essential for virulence in non-flowering plant tissues. In addition to its role in promoting necrosis, syringomycin exhibits biosurfactant properties that facilitate pathogen spread^37^, which we observed in wound-based infection assays in liverwort thalli. Collectively, our data demonstrate that syringomycin is crucial for disease establishment in diverse plant hosts. Future studies are required to understand how toxins like syringomycin contribute to virulence alongside effectors, particularly those interfering with processes at the plasma membrane.

While phytopathogenic *Pseudomonas* species deploy cyclic lipopeptides to promote infection in plants, these toxins also have profound impacts in fungi, oomycetes, algae, and insects. The lipopeptides massetolide A, viscosin, and WL-3 impair the viability of oomycete (*Phytophthora* and *Pythium*) zoospores or hyphae^40,41^, similar to our observation of SRE inhibiting *P. infestans*. Moreover, fuscopeptin and syringotoxin produced by the sheath rot pathogen *P. fuscovaginae* participate in disease establishment of rice panicles while also exhibiting antifungal activity against the plant pathogen *Rhizoctonia solani*^42^. Plant-beneficial *Pseudomonas* species like *P. protegens* encode toxins with both antimicrobial and anti-insect activity, like the cyclic lipopeptide orfamide A that exerts insecticidal activity in *Plutella xylostella* and *Drosophila melanogaster*^43,44^. Orfamide A production by *P. protegens* also disrupts calcium homeostasis in the green alga *Chlamydomonas reinhardtii,* which results in algal deflagellation and immobilization^45^. Intriguingly, *Mycetocola lacteus* helper bacteria can protect algae from deflagellation by linearizing orfamide A, which allows *M. lacteus* helper bacteria to access *C. reinhardtii* micronutrients^46^. Together these works highlight the ecological importance of cyclic lipopeptide toxins for diverse host-microbe interactions in the soil as well as the phyllosphere.

Our work demonstrates that the *Pseudomonas syringae* species complex infects a diverse range of host plants that includes liverworts and ferns alongside flowering plants. In particular, we highlight the role of the lipopeptide phytotoxin syringomycin, which works alongside type-3 effectors to establish disease. While syringomycin was a key factor underpinning the broad host virulence of PG2 isolates, we also observed susceptible interactions between non-flowering plants and related *Pseudomonas* species like *P. cichorii, P. asplenii,* or *P. viridiflava.* This hints to a wide spectrum of virulence mechanisms that may enable plant infection. Future studies aimed at resolving bacterial virulence across divergent systems may yet reveal fundamental vulnerabilities that microbes converge upon to cause disease.

## ACKNOWLEDGEMENTS

We thank all researchers who donated *Pseudomonas* isolates to stock centers (ICMP, LMG). In particular, we also thank Rose Williams (Manaaki Whenua – Landcare Research, New Zealand) for organizing material transfers and helpful post-order follow up. We thank Andy Plackett (University of Birmingham, UK) for providing *C. richardii* spores and for advice on fern growth and maintenance. We thank Jon Takemoto (University of Utah, United States) for invaluable advice on syringomycin purification. We thank Steven E. Lindow (University of California, Berkeley, United States) for providing wildtype B728a and the ΔsyrP mutant, Renier van der Hoorn (University of Oxford, United Kingdom) for providing the B728a ΔhrcC mutant, and Hirofumi Nakagami (Max Planck Institute for Plant Breeding, Germany) for providing the PtoDC3000-lux strain. The *P. syringae* bioluminescence vectors pBJ2 (Addgene plasmid # 167131) and pBJ4 (Addgene plasmid # 167133) were gifts from Akira Mine (Kyoto University, Japan).

## AUTHOR CONTRIBUTIONS

PC designed research; KG, KSC, ET, RA, JR, and PC performed research; KG, KSC, AST, RA, MR, MV, JFW, and PC analyzed data; AST, MV, JFW, and PC performed bioinformatic analyses including RNA-sequencing and orthology analysis; MR performed analytical chemistry and purified syringomycin; PC prepared figures and final datasets; PC wrote the paper with contributions from all authors.

## DATA AVAILABILITY

All relevant gene identifiers and dataset accession numbers are provided in the manuscript.

## MATERIALS & METHODS

### Plant growth details

The liverwort *Marchantia polymorpha* (Tak-1; Takaragaike-1) was axenically cultivated by passaging asexual gemmae ontop of a sterile nylon mesh (100 micron, Normesh) placed on one-half–strength MS-B5 (Murashige and Skoog with B5 vitamins) media (pH 6.7) under long-day photoperiod (16 hours of light; ∼80 uE light intensity) at 20-22 °C. The fern *Ceratopteris richardii* (Hn-n) was grown from surface sterilized spores (15% v/v commercial bleach and 0.1% Tween-20 for 10 minutes, followed by 5 washes with distilled water) plated onto C-fern growth media^47^ and grown at 25 °C with a long-day photoperiod (16 hours light; ∼80-120 uE) for 4-6 weeks until sporophytes developed. Sporophytes with at least 2 fronds were transplanted to soil and grown under controlled conditions with a long-day growth period (16 hours of light; ∼120-150 μE) at 22 °C and ∼80% relative humidity. *Nicotiana benthamiana* were grown in soil under controlled conditions with a temperature of 22 °C and a long day photoperiod (16 hours of light; ∼120-150 μE light intensity).

### Bacterial growth and plant infection assays

All *Pseudomonas* isolates (listed in Data S1) were grown axenically in King’s B (KB) media at 28 °C with appropriate antibiotics until the exponential phase. Bacteria were collected by centrifugation, resuspended in 5 mM MgCl_2_ and adjusted to 1×10^6^ colony forming units (CFU) ml^−1^. Bacteria were pressure infiltrated into fully expanded leaves or fronds of 4-week-old *N. benthamiana* or *C. richardii,* respectively, using a 1 ml needles syringae. For *Marchantia,* bacterial suspensions were vacuum infiltrated into 4-week-old thalli collected from nylon mesh growth plates and then placed into a fresh petri dish containing wetted sterile Whatman paper and returned to initial growth conditions. To quantify bacterial growth *in planta,* three sets containing 8 tissue discs (4 mm diameter) were collected and placed in fresh 10 mM MgCl_2_ containing 0.1% Silwet L-77 before being shaken at 170 rpm for 1 hour. Serial dilutions prepared in 10 mM MgCl_2_ were subsequently plated on KB media, incubated at 28 °C, and counted 1-2 days later.

### RNA isolation, cDNA synthesis, and qRT-PCR analyses

Plant total RNA was collected from flash frozen tissue using the Spectrum Plant RNA Extraction Kit (Protocol A) with on-column DNAse treatment following the manufacturer’s instructions (Sigma). cDNA was then synthesized from 2 µg of total RNA using Superscript II Reverse Transcriptase (Invitrogen) following the manufacturer’s instructions. Bacterial total RNA was collected from overnight cultures (KB or MM) or isolated from plant tissues using established protocols^48^. cDNA was then synthesized from 800 ng of total RNA using SuperScript IV VILO Master Mix (Invitrogen) following the manufacturer’s instructions. Diluted plant or bacterial cDNA (10x) was used for qRT-PCR reactions with Roche SYBR mix and gene-specific primers (Table S1) as previously described^26^. Expression levels were normalized to internal controls. All statistical analyses (ANOVA, Tukey’s HSD) were performed in R and graphs were generated in GraphPad Prism (v10.1.0).

### Library preparation and RNA-sequencing

mRNA from mock-inoculated (MgCl_2_) or *P. syringae* pv. *syringae (*B728a^WT^, ΔhrcC, or Δsyrp) infected *M. polymorpha*, *C. richardii,* and *N. benthamiana* plants at 24 hours post treatment were purified from DNAse-treated total RNA (as described above) using Poly(A) selection and were then fragmented. Library preparation of cDNA was performed using the TruSeq® RNA Sample Preparation Kit (Illumina, US) according to manufacturer’s instructions. Sequencing was performed on the Illumina NovaSeq X in 150 paired end mode. De-multiplexed samples were used for downstream expression analyses. Raw fastq data are accessible at http://www.ncbi.nlm.nih.gov/sra/ under the accession number PRJNA1126102.

### RNA-sequencing read processing and differential gene expression analysis

The program fastqc v.0.11.9 (https://www.bioinformatics.babraham.ac.uk/projects/fastqc) was used to assess the quality of the raw reads. The Kakapo v.0.9.0 pipeline^49^ was then used to process and clean the sequencing reads. In short, Kakapo first uses Rcorrector v.1.0.5^50^ to correct random base pair sequencing errors. Trimmomatic v.0.39^51^ was used to remove adapter sequences, low quality reads and bases with low Phred scores. Bowtie2 v.2.4.4^52^ was used to filter out sequences of interest, in this case, *Pseudomonas* reads (CP074578.1 and CP074579.1) and Kraken2 v2.1.2^53^, with the databases provided by Kakapo, was used to filter out further contaminants. Once processed, fastqc was used to ensure the cleaned and filtered reads had increased quality.

The abundances of the corrected reads were quantified with Kallisto v.46.2^54^ using their respective species coding sequences, *Nicotiana benthamiana* (NbLAB; https://bioweb01.qut.edu.au/benthTPM/download.html)^55^, *Ceratopteris richardii* (v2.1; https://phytozome-next.jgi.doe.gov/info/Crichardii_v2_1)^56^, and *Marchantia polymorpha* (MpTak1_v5.1; https://marchantia.info/about_genome/)^57^ as references. Two approaches were taken for the differential gene expression analysis. One was a pairwise approach using all the data and the other was a multi-treatment comparison using further data filtering. In both cases, the abundances were converted into counts using the R package tximport^58^. DESeq2^59^ was used to identify differentially expressed genes in a pairwise manner. For the multi-treatment approach that was used for hierarchical clustering and analysis of co-variance, any genes that had less than 10 reads mapping across all samples were removed prior to the DESeq2 analysis. The differentially expressed genes were those with an adjusted p-value of less than 10^−3^ and a log_2_ fold change greater than or equal to the absolute value 1.5. Heatmaps of differentially expressed genes were generated for each species using the R package pheatmap (https://www.rdocumentation.org/packages/pheatmap/versions/1.0.12), with variance stabilized counts. To compare the infection transcriptomes of *M. polymorpha, C. richardii,* and *N. benthamiana*, we used previously defined sets of orthologous protein groups (orthogroups)^26^ generated using OrthoFinder^25^. The MBEX online tool (https://marchantia.info/mbex/)^60^ was used to perform functional enrichment analyses with a significance cutoff of FDR (False Discovery Rate) ≤ 0.05.

### Syringomycin purification and application experiments

Syringomycin E (SRE) was purified from *P. syringae* pv. *syringae* B301D grown in freshly prepared potato dextrose broth at 25 °C for 10 days following an adapted protocol^61,62^ (see the supplementary information for the full protocol). This was facilitated by screening for bioactive SRE-containing fractions capable of suppressing *Phytophthora infestans* (88069) hyphal growth on PDA plates grown at 18 °C for at least 6 days. Purified SRE was aliquoted, freeze-dried, and stored at −20 °C. For SRE application experiments, SRE aliquots were resuspended in sterile water to a stock concentration of 0.5 μg μl^−1^, with all subsequent dilutions being made in water. Activity assays in *Nicotiana* and *Ceratopteris* were performed by pressure infiltrating SRE into fully expanded leaves/fronds of 4-week-old plants using a needless syringe. SRE assays in *Marchantia* were performed by spotting 10 µL droplets onto the ventral surface of 4-week-old thalli, or by vacuum infiltrating 4-week-old thalli grown on nylon mesh in one-half-strength MS-B5 media. SRE impact on plant tissues was assessed 24 hours post treatment.

To perform SRE rescue assays, we first transformed bioluminescence reporter constructs (pBJ4 or pBJ2) into *P. syringae* following previously described electroporation and selection protocols^19^. Since purified SRE stocks are limited, we developed a localized assay whereby sterile toothpicks dipped into freshly grown bacterial colonies (on KB media) were delivered into thalli by gentle wounding. Immediately following delivery, we applied 5 µL droplets of either water (mock) or SRE (0.5 μg μl^−1^) into the wound site. Luminescence was detected 2 days later by NightOwl imaging, or by collecting 4 mm tissue discs around the inoculation site (minimum 40 discs from 10 plants per treatment), floating them in 100 µL water in a 96 well plate, and measuring luminescence glow using a Varioskan plate reader.

### Statistical Analyses

Details regarding statistics analyses performed for a given experiment are outlined in the relevant figure legends. Here, the statistical tests used, the sample size (i.e., n = number of independently infected plants) and dispersion/precision measurements are also provided (error bars, p value cutoffs, etc.). Statistical analyses for transcriptomics are described in the Methods above. Statistical analysis of qRT-PCR data were performed in R software and are described in appropriate figure legends.

## SUPPLEMENTARY INFORMATION

**Figure S1. Comparative analysis of plant pathogenic *P. syringae* isolates in diverse hosts**

**Figure S2. Transcriptome-wide responses to *P. syringae* infection across diverse plants.**

**Figure S3. Purification and bioactivity of Syringomycin E**

**Figure S4. Syringomycin promotes necrotic cell death in plants**

**Figure S5. Syringomycin enhances *P. syringae* growth in *Marchantia***

**Data S1. *Pseudomonas* isolates and infection-related information**

**Data S2. Plant infection phenotypes**

**Data S3. RNA-sequencing analysis of diverse hosts infected with *P. syringae* B728a**

**Table S1. Primers used in this study**

**Supplementary Method – Extended SRE Purification Protocol**

