## Supplementary material for "A Necrotizing Toxin Promotes *Pseudomonas syringae* Infection Across Evolutionarily Divergent Plant Lineages": Data S2.pdf

**Dataset S2. Plant infection phenotypes**

*Marchantia* *Ceratopteris* *Nicotiana*

PG1:

Pat LMG5057

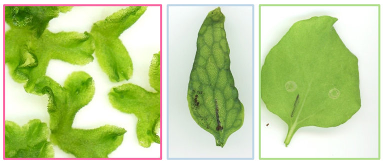

Pdp ICMP529

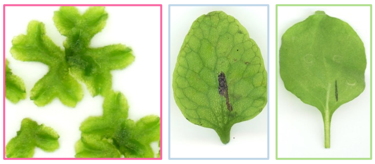

Psp ICMP16929

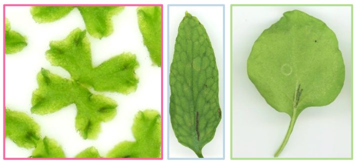

Pto DC300

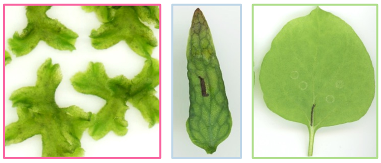

Pto ICMP7230

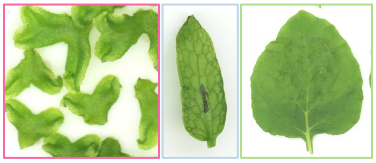

PG2:

Paf LMG5095

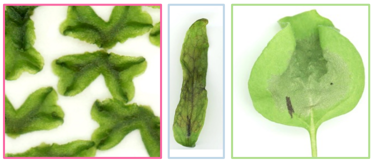

Paf ICMP1852

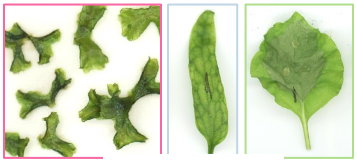

Paf ICMP5011

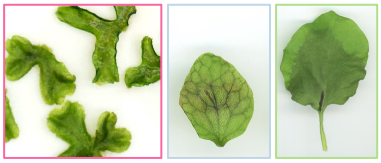

Pcr ICMP17001

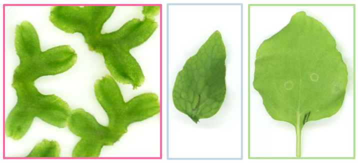

Per ICMP4967

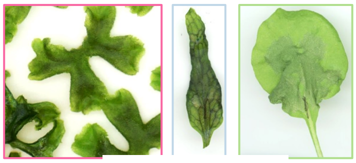

Plp LMG2206

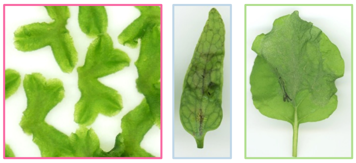

Ppi LMG5079

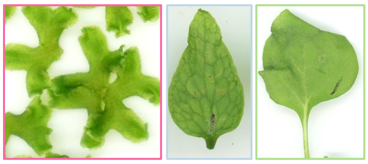

Ppp ICMP4048

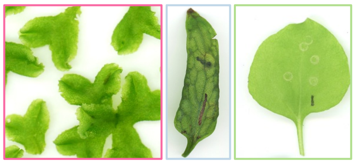

Psy ICMP4917

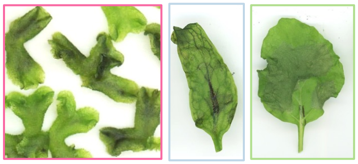

Psy B728a

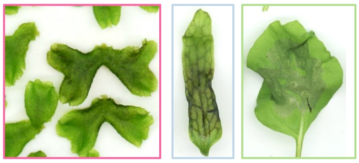

Ptt LMG5059

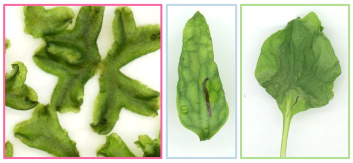

Ptt ICMP11935

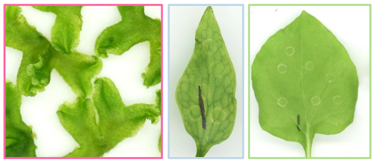

Ptt ICMP4388

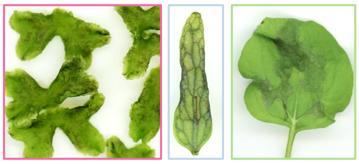

PG3:

Psa ICMP19198

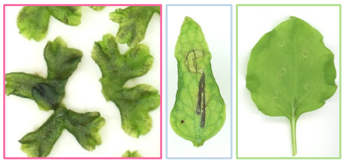

Pcs ICMP9419

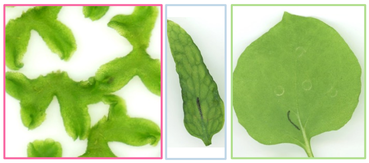

Pcu ICMP11894

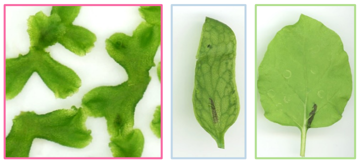

Pda ICMP9757

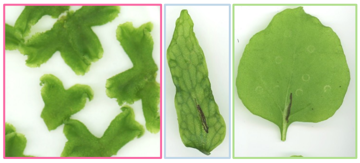

Phi ICMP9623

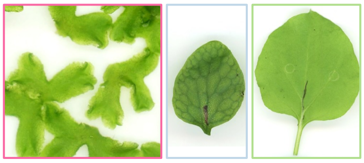

Pho ICMP7840

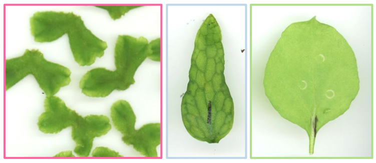

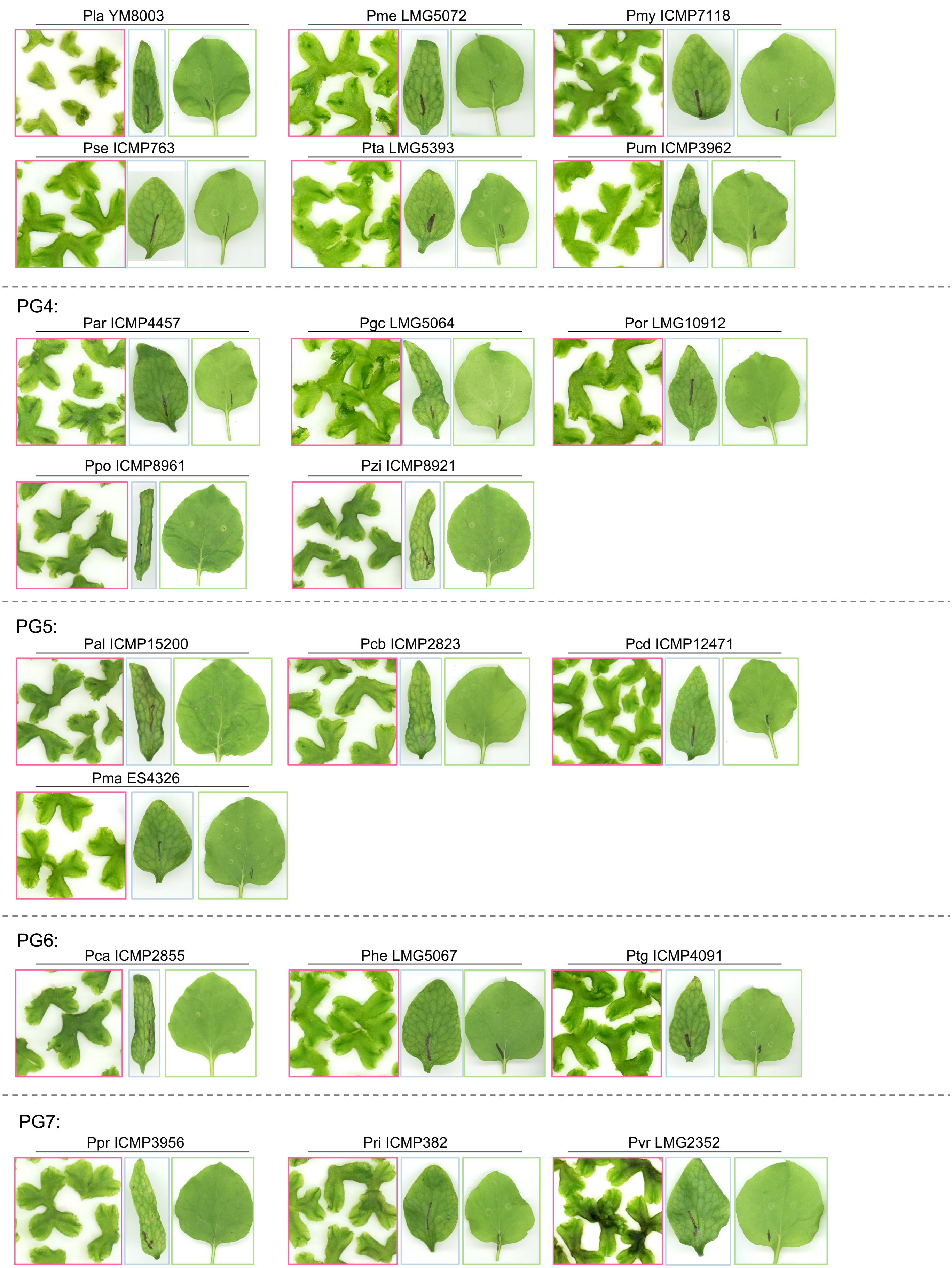

Pla YM8003
Pme LMG5072
Pmy ICMP7118
Pse ICMP763
Pta LMG5393
Pum ICMP3962
PG4:
Par ICMP4457
Pgc LMG5064
Por LMG10912
Ppo ICMP8961
Pzi ICMP8921
PG5:
Pal ICMP15200
Pcb ICMP2823
Pcd ICMP12471
Pma ES4326
PG6:
Pca ICMP2855
Phe LMG5067
Ptg ICMP4091
PG7:
Ppr ICMP3956
Pri ICMP382
Pvr LMG2352

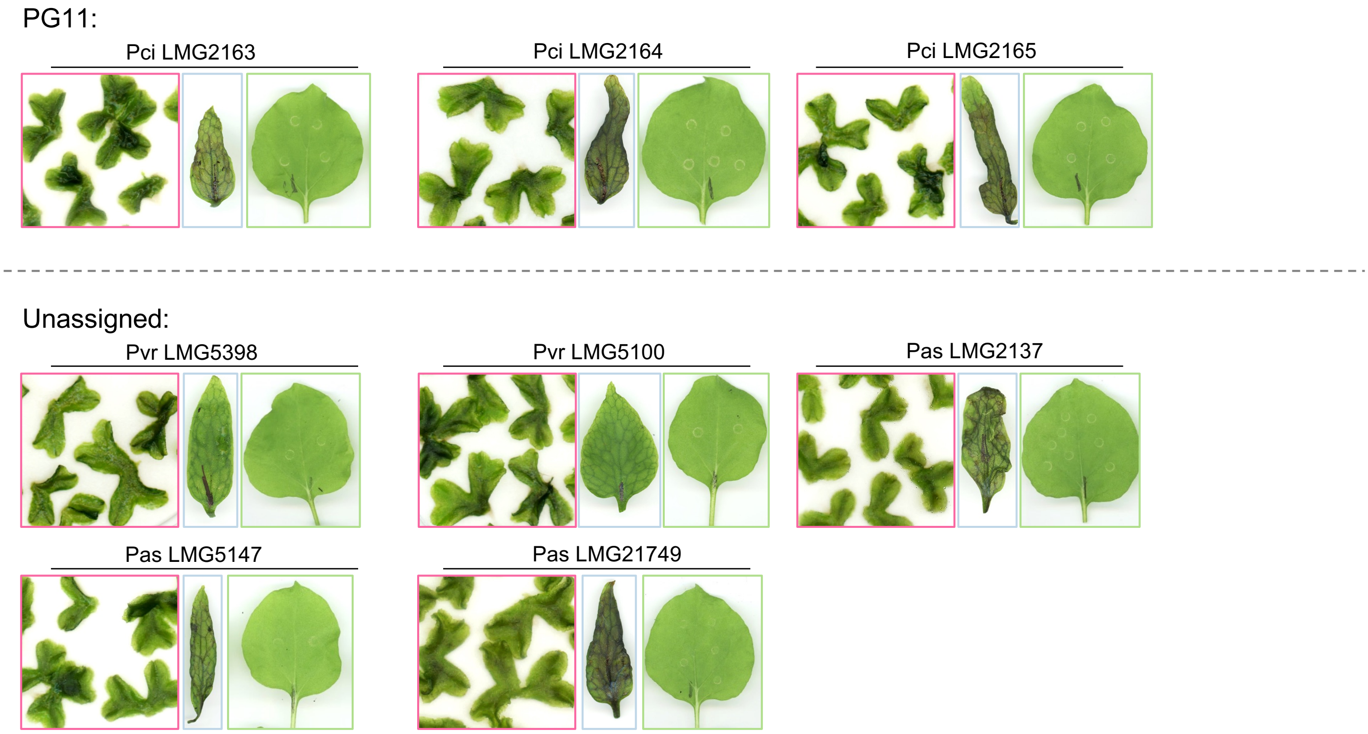


**Data S2. Plant infection phenotypes**

Representative infection phenotypes of *Marchantia polymorpha, Ceratopteris richardii, and Nicotiana benthamiana* infected with diverse *Pseudomonas* isolates (related to Figure 1). Images are organized by *P. syringae* species complex phylogroup (PG) with appropriate isolate identifiers labelled above each set of plants (see Data S1 for isolate information). Images were taken 3 days post infection (dpi) for *M. polymorpha* and *C. richardii,* or 2 dpi for *N. benthamiana.* Tissue from at least 8-12 individual plants was harvested for each treatment/replicate. Infections were performed at least twice with similar results.
