## Supplementary material for "A Necrotizing Toxin Promotes *Pseudomonas syringae* Infection Across Evolutionarily Divergent Plant Lineages": Figure S1.pdf

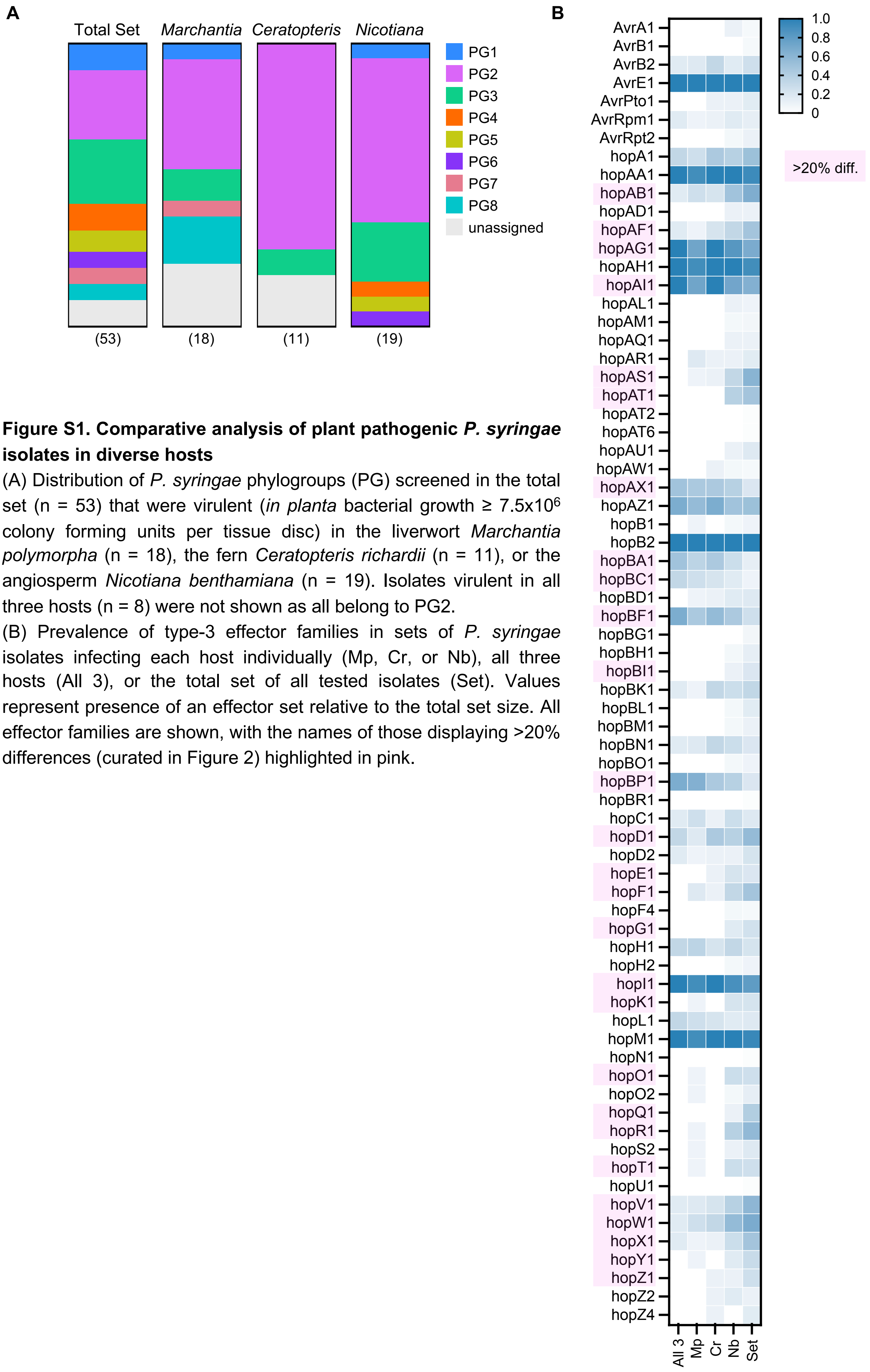


**Figure S1. Comparative analysis of plant pathogenic *P. syringae* isolates in diverse hosts**

(A) Distribution of *P. syringae* phylogroups (PG) screened in the total set (n = 53) that were virulent (*in planta* bacterial growth ≥ $7.5 \times 10^6$ colony forming units per tissue disc) in the liverwort *Marchantia polymorpha* (n = 18), the fern *Ceratopteris richardii* (n = 11), or the angiosperm *Nicotiana benthamiana* (n = 19). Isolates virulent in all three hosts (n = 8) were not shown as all belong to PG2.

(B) Prevalence of type-3 effector families in sets of *P. syringae* isolates infecting each host individually (Mp, Cr, or Nb), all three hosts (All 3), or the total set of all tested isolates (Set). Values represent presence of an effector set relative to the total set size. All effector families are shown, with the names of those displaying >20% differences (curated in Figure 2) highlighted in pink.
