## Supplementary material for "A Necrotizing Toxin Promotes *Pseudomonas syringae* Infection Across Evolutionarily Divergent Plant Lineages": Figure S2.pdf

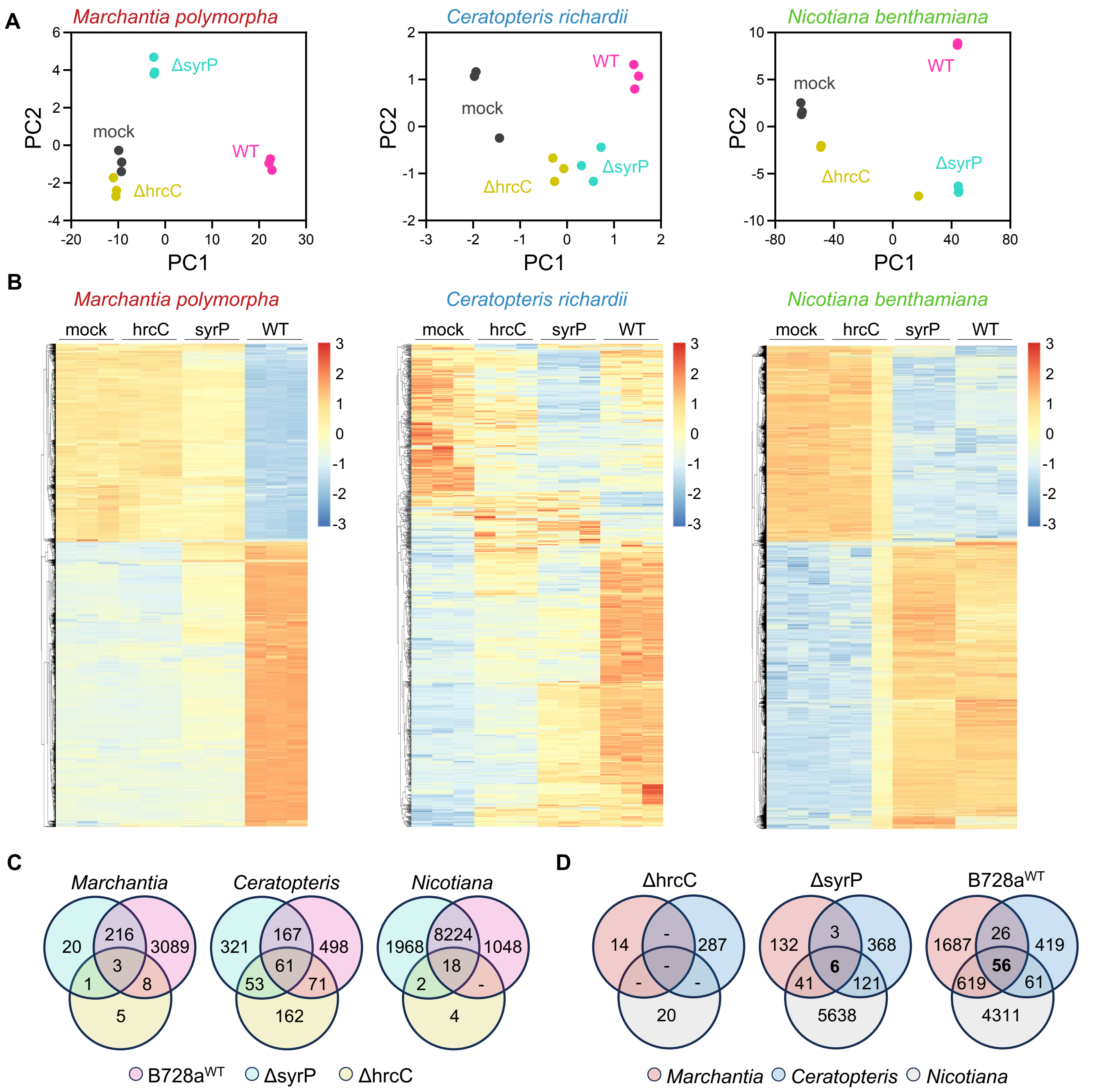


**Figure S2. Transcriptome-wide responses to *P. syringae* infection across diverse plants.**

(A) Principal component analysis (PCA) of *Marchantia, Ceratopteris*, and *Nicotiana* transcriptome replicates. Three samples per replicate of host tissues collected 24 hours post infiltration (hpi) with mock (10 mM $MgCl_2$), wildtype *P. syringae* pv. *syringae* B728a (B728a$^{WT}$), ΔhrcC, or ΔsyrP mutants are shown.

(B) Hierarchical clustering of significant differentially expressed genes (DEGs) in each host in response to mock-inoculation or *P. syringae* infection (adjusted P-value < $10^{-3}$, log fold change (|LFC| ⩾ 1.5).

(C) Venn diagram showing overlapping sets of total DEGs between infection treatments in a given species.

(D) Venn diagrams of showing overlapping sets of conserved differentially expressed orthologous genes that are shared between species for a given *P. syringae* treatment.
