## Supplementary material for "A Necrotizing Toxin Promotes *Pseudomonas syringae* Infection Across Evolutionarily Divergent Plant Lineages": Figure S3.pdf

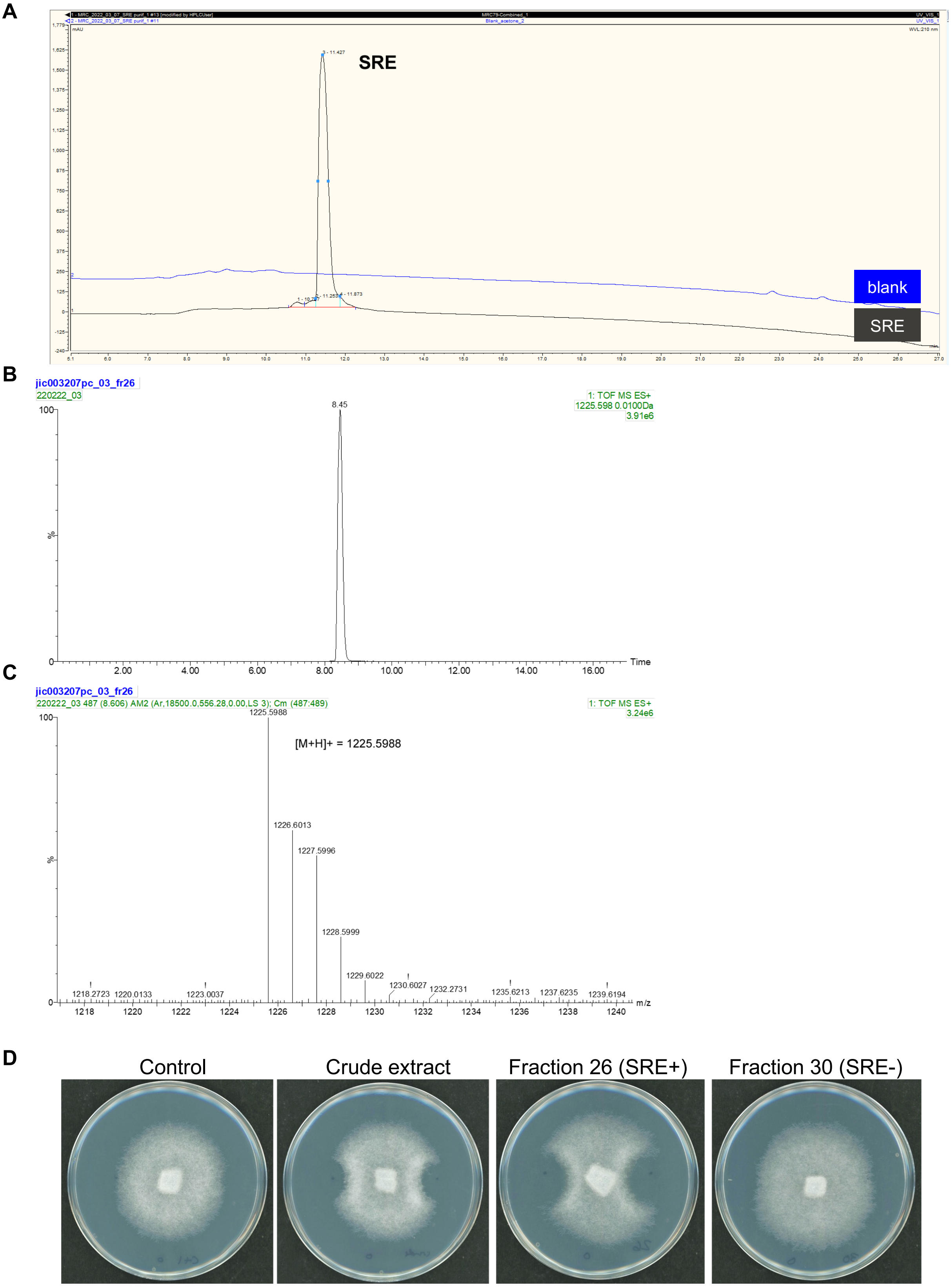


**Figure S3. Purification and bioactivity of Syringomycin E**

**(A)** Purity check of purified Syringomycin E (SRE) by HPLC comparing a ‘blank’ control versus SRE. Detection by UV at 210 nm. Syringomycin E elutes at 11.4 minutes. Purity 94%.

**(B)** HR-MS of purified SRE on Synapt G2-Si: extracted ion chromatogram for m/z 1225.6.

**(C)** HR-MS of purified SRE on Synapt G2-Si: pseudomolecular ion for $[M+H]^+$ = $C_{53}H_{86}ClN_{14}O_{17}$ calculated 1225.5980, observed 1225.5988, error 0.65 ppm.

**(D)** Anti-oomycete activity of SRE. 6-day-old PDA plates pre-spotted with a buffer control, total crude SRE extract, an SRE-containing fraction (26), or a fraction lacking SRE were inoculated with a central hyphal plug of *Phytophthora infestans* (88069). Two plates per treatment were assayed in two independent experiments.
