## Supplementary material for "A Necrotizing Toxin Promotes *Pseudomonas syringae* Infection Across Evolutionarily Divergent Plant Lineages": Figure S4.pdf

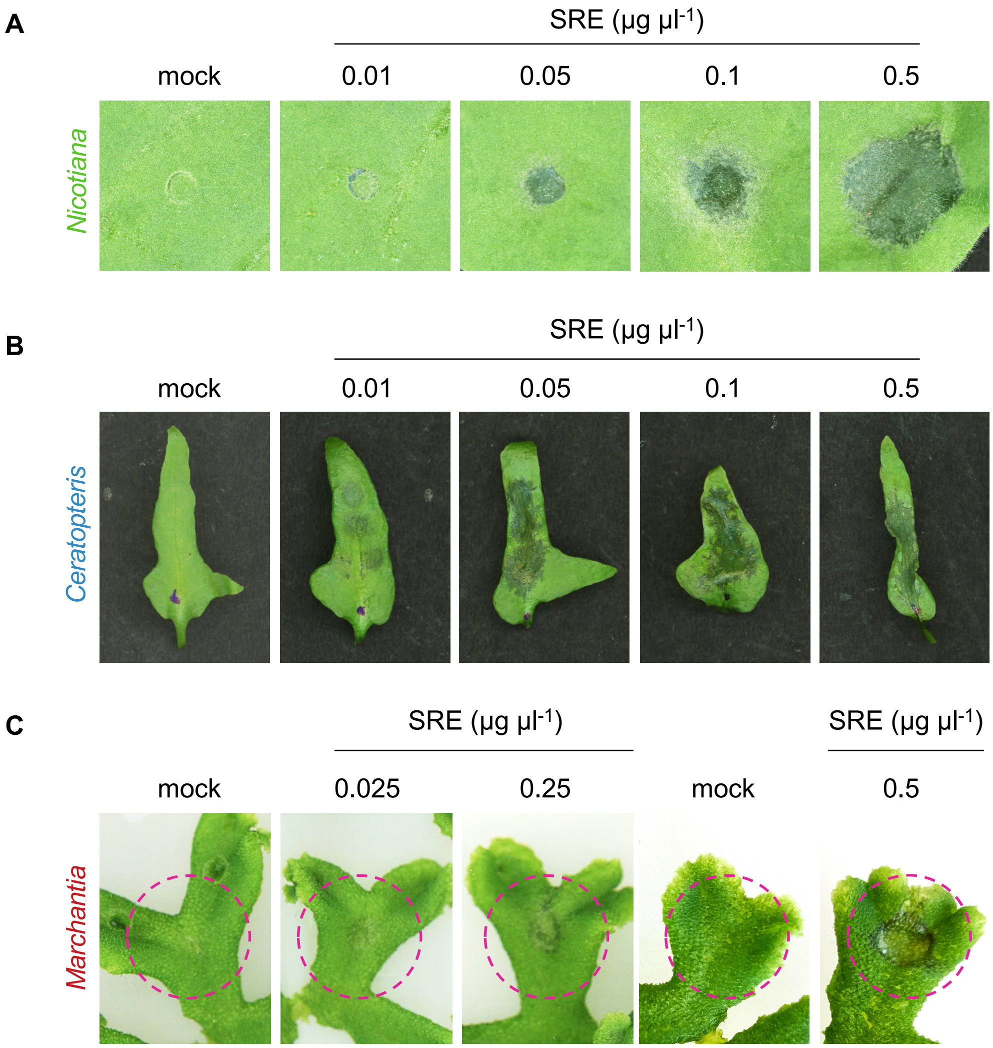


**Figure S4. Syringomycin promotes necrotic cell death in plants**

(A) Dose-response of *Nicotiana benthamiana* leaves to infiltration with indicated concentrations of purified syringomycin E (SRE) relative to a mock-treated (distilled water) control (n = 6 plants per treatment). Images were taken 24 hours post infiltration (hpi). This experiment was performed three times with similar results.

(B) Dose-response of *Ceratopteris richardii* fronds to infiltration with indicated concentrations of purified SRE relative to a mock-treated (distilled water) control (n = 8). Images were taken 24 hpi. This experiment was performed three times with similar results.

(C) Dose-response of *Marchantia polymorpha* thalli to droplet application (5 µl) of the indicated concentrations of purified SRE relative to a mock-treated (distilled water) control (n = 10). Images were taken 24 hours post treatment. This experiment was performed at least three times with similar results.
