## Supplementary material for "A Necrotizing Toxin Promotes *Pseudomonas syringae* Infection Across Evolutionarily Divergent Plant Lineages": Figure S5.pdf

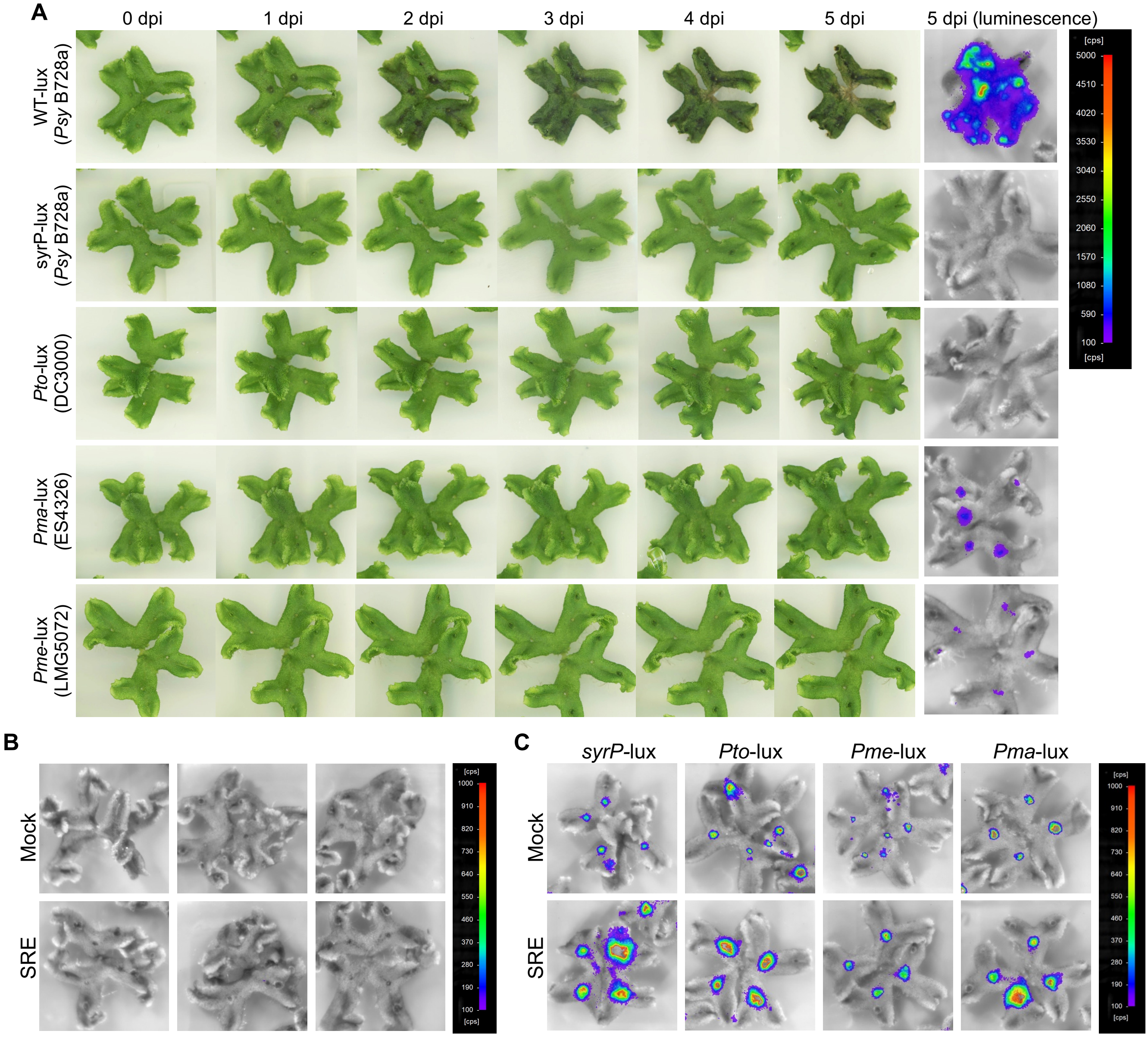


**Figure S5. Syringomycin enhances *P. syringae* growth in *Marchantia***

(A) Time course infection assays of 4-week-old *Marchantia polymorpha* thalli wound-inoculated with bioluminescent reporter isolates *P. syringae* pv. *syringae* B728a (WT-lux), the B728a ΔsyrP-lux mutant, *P. syringae* pv. *tomato* DC3000 *(Pto-lux), P. syringae* pv. *maculicola* ES4326 *(Pma-lux*), or *P. syringae* pv. *mellea* LMG5072 *(Pme-lux*). Images show representative infection phenotypes at the indicated days post inoculation (dpi) alongside bioluminescence measurements taken at 5 dpi (n = 8). Values above or below the indicated scale are not false-colored.

(B) Control assay showing the absence of luminescence detection for mock (distilled water) or ectopic syringomycin E (SRE, 0.5 µg $\mu l^{-1}$) application of *Marchantia* thalli (n = 8). Values above or below the indicated scale are not false-colored.

(C) Luminescence detection of wound-inoculated *P. syringae* reporter isolates treated with distilled water (Mock) or SRE (0.5 µg $\mu l^{-1}$). Images are taken at 2 dpi (n = 10). Full quantification data is provided in Figure 4. Values above or below the indicated scale are not false-colored.
