## Supplementary material for "A Necrotizing Toxin Promotes *Pseudomonas syringae* Infection Across Evolutionarily Divergent Plant Lineages": MarchantiaCode.R

```
library( "DESeq2" )
library("rhdf5")
library("tximport")
library("pheatmap")

#Set the directory to where the expression analysis output is
setwd("/Users/Walker/Dropbox/LabProjects/PhilProject_2/PCA_And_HeatMap/ForPhil/Abundances/")

#make the directories, each one of these is in the path and have the abundance data
dirs <- c("hdm-1_mapping","hdm-2_mapping","hdm-3_mapping","hdh-1_mapping","hdh-2_mapping","hdh-3_mapping","hds-1_mapping","hds-2_mapping","hds-3_mapping","hdw-1_mapping","hdw-2_mapping","hdw-3_mapping")

#input the files, this gets the path and stores the file names as files
files <- file.path(getwd(),dirs,"abundance.tsv")
names(files) <- paste0(txIdCol = c("Mock1","Mock2","Mock3","HRCC1","HRCC2","HRCC3","SYRP1","SYRP2","SYRP3","WILDTYPE1","WILDTYPE2","WILDTYPE3"))

#Kallisto is a pseudomapper so the data needs to be converted using tximport
txi.kallisto <- tximport(files, type = "kallisto", txOut = TRUE)

#Convert the tximport data to count data for downstream analysis
countdata <- as.matrix(txi.kallisto)

#Build the DESeq dataframe
sampleTable <- data.frame(
  condition = factor(c(
    rep("Mock", 3),
    rep("HRCC", 3),
    rep("SYRP", 3),
    rep("WILDTYPE", 3))))
sampleTable$condition <- relevel(coldata$condition, ref = "Mock")

#convert the row names for DESeq2
rownames(sampleTable) <- colnames(txi.kallisto$counts)

#Change to DESeq2 dataframe
dds <- DESeqDataSetFromTximport(txi.kallisto, sampleTable, design=~condition)

#remove rows that have >= 10 reads in total, more strict filtering is applied via independent filtering on the mean of normalised counts within results function
dds <- dds[ rowSums(counts(dds)) > 10, ]


#Start DESeq2
dds <- DESeq(dds)

#Get the results names (Check the right comps are being made)
resultsNames(dds)

#do all pairwise comps
res_mock_hrcc <- results(dds, c("condition", "Mock", "HRCC"), alpha = 0.001)
res_mock_wildtype <- results(dds, c("condition", "Mock", "WILDTYPE"), alpha = 0.001)
res_mock_syrp <- results(dds, c("condition", "Mock", "SYRP"), alpha = 0.001)

#order each by the p-value
res_ordered_mock_hrcc <- res_mock_hrcc[order(res_mock_hrcc$padj),]
res_ordered_mock_wildtype <- res_mock_wildtype[order(res_mock_wildtype$padj),]
res_ordered_mock_syrp <- res_mock_syrp[order(res_mock_syrp$padj),]

#Identify number of p-values less that 0.001
sum(res_mock_hrcc$padj < 0.001, na.rm=TRUE)
sum(res_mock_wildtype$padj < 0.001, na.rm=TRUE)
sum(res_mock_syrp$padj < 0.001, na.rm=TRUE)

#Get the significant genes in each comparison
#resSig_mock_hrcc <- as.data.frame(subset(res_mock_hrcc, padj < 0.001 & abs(log2FoldChange) >= 2))
#resSig_mock_hrcc <- subset(res_mock_hrcc, padj < 0.001 & abs(log2FoldChange) >= 2)

resSig_mock_hrcc <- as.data.frame(subset(res_mock_hrcc, padj < 0.001 & abs(log2FoldChange) >= 1.5))
resSig_mock_hrcc <- subset(res_mock_hrcc, padj < 0.001 & abs(log2FoldChange) >= 1.5)

#resSig_mock_wildtype <- as.data.frame(subset(res_mock_wildtype, padj < 0.001 & abs(log2FoldChange) >= 2))
#resSig_mock_wildtype <- subset(res_mock_wildtype, padj < 0.001 & abs(log2FoldChange) >= 2)

resSig_mock_wildtype <- as.data.frame(subset(res_mock_wildtype, padj < 0.001 & abs(log2FoldChange) >= 1.5))
resSig_mock_wildtype <- subset(res_mock_wildtype, padj < 0.001 & abs(log2FoldChange) >= 1.5)

#resSig_mock_syrp <- as.data.frame(subset(res_mock_syrp, padj < 0.001 & abs(log2FoldChange) >= 2))
#resSig_mock_syrp <- subset(res_mock_syrp, padj < 0.001 & abs(log2FoldChange) >= 2)

resSig_mock_syrp <- as.data.frame(subset(res_mock_syrp, padj < 0.001 & abs(log2FoldChange) >= 1.5))
resSig_mock_syrp <- subset(res_mock_syrp, padj < 0.001 & abs(log2FoldChange) >= 1.5)

#### heatmaps of the most significant DEG, overview

vsd <- vst(dds, blind = FALSE) # variance-stabilised counts
head(assay(vsd),3)


select <- order(rowMeans(counts(dds,normalized=TRUE)),decreasing=TRUE)
heatmap_matrix <- assay(vsd)[select,]
rows_in_matrix <- row.names(heatmap_matrix)

#combine the significant genes into one matrix, and make sure it's a unique list
rows_to_keep <- unique(c(row.names(resSig_mock_hrcc),row.names(resSig_mock_wildtype),row.names(resSig_mock_syrp)))
rows_for_input <- match(rows_to_keep, rows_in_matrix)
final.matrix <- heatmap_matrix[rows_for_input, ]
#length(rows_for_input)

#generate a gene heatmap
tiff("Marchantia_gene_heat_unclustered.tiff", 1000, 1000, pointsize=20)
pheatmap(final.matrix,show_rownames=FALSE,show_colnames = TRUE,scale="row",cluster_cols = FALSE)
#tiff("Marchantia_gene_heat.tiff", 1000, 1000, pointsize=20)
#pheatmap(final.matrix,show_rownames=FALSE,show_colnames = TRUE,scale="row")
dev.off()

#generate a pca
png("Marchantia_qc-pca.png", 1000, 1000, pointsize=20)
plotPCA(vsd[rows_for_input], intgroup="condition")
dev.off()

#Write the PC values to a csv
write.csv(plotPCA(vsd[rows_for_input], intgroup="condition",returnData=TRUE),"Marchantia_PC_Values.csv")
```
