## Supplementary material for "A Necrotizing Toxin Promotes *Pseudomonas syringae* Infection Across Evolutionarily Divergent Plant Lineages": Supplementary Method - Extended SRE Purification Protocol.pdf

### SUPPLEMENTARY METHODS

#### Detailed Syringomycin E purification protocol

Our method of Syringomycin E purification was adapted from Zhang and Takemoto (1987). In brief, Syringomycin E was extracted and purified from *Pseudomonas syringae* pv. *syringae* strain B301D grown in potato dextrose broth (PDB). It is critical that PDB be made fresh rather than using commercial mixes. First, 800 g of cubed potatoes was boiled in 3 L of distilled water for 30 minutes, then when cooled the solution was filtered through 2 layers of cheese cloth. Next, glucose (80 g, 2 w/v%) and casamino acids (16 g, 0.4 w/v%) were added and the resulting mixture was adjusted to a final volume of 4 L and stirred with a magnetic stirring bar until dissolved. The medium was dispensed into 500 ml culture flasks (10 flasks) and sterilised by autoclaving. Starter cultures of *P. syringae* B301D were grown by shaking (180-200 rpm) at 28 $^{o}$C from freshly streaked bacteria grown on KB media. For syringomycin purification, eight flasks (4 L) were inoculated with *P. s.* pv. *syringae* B301D (1 ml) and were allowed to grow as standing cultures at 24 $^{o}$C for 10 days. Two flasks (1 L) of uninoculated control media were separated and allowed to stand under the same conditions. These were used as unspent media control.

The flasks containing either culture or the unspent media were chilled to 4 $^{o}$C and acidified acetone (4 ml HCl per 1 l liter acetone) was added to give a 1:1 mixture media/acetone. The flasks were allowed to gently stir overnight at 4 $^{o}$C. Cellular debris were removed by centrifugation (9000 g, 20 min, 4 $^{o}$C) (Sorvall Lynx 4000). The supernatant was concentrated to 200 ml *in vacuo* by rotary evaporation with the evaporator water bath temperature set to 45 $^{o}$C. Acetone (300 ml, 60% v/v acetone solution) was added and the suspension was allowed to stir gently overnight at 4 $^{o}$C. However, the best results were achieved when left for 3 days.The suspension was centrifuged (15123 g, 30 min, 4 $^{o}$C, Sorvall Lynx 4000) and acetone was removed by rotary evaporation. The residue was filtered through filter paper (Buchner funnel and water aspirator vacuum was required) and applied on a RP-C18 column (SNAP KP-C18-HS, 120 g, Biotage). The target compound was eluted using gradient of buffer B (i-PrOH + 0.1% TFA) against buffer A ($H_2O$ + 0.1% TFA) at flow rate 30 ml/min. First 0% B for 12 minutes (360 ml), then gradient to 100 % B over 18 minutes (540 ml), hold for 6 minutes (180 ml) with monitoring UV at 254 nm. The fractions from Biotage were evaporated to dryness (Genevac EZ-2 Elite, HPLC setting, 30 $^{o}$C, 24 h).

The residue was dissolved in $H_2O$/acetone 4:6 (3 ml) and each fraction was evaluated for presence of Syringomycin E using HPLC (Agilent 1260) on a XB-C18 column (100 Å, 100 x 4.6 mm, 5 µm) using gradient of B (MeCN + 0.1% FA) against A ($H_2O$ + 0.1% FA). The method started at 10% B for 1 min, then gradient to 98% over 10 min, hold for 2 min, back to 10 % over 0.1 min and hold 1.9 min, at flow rate 1 ml/min with detection by UV (210, 230, 250 and 400 nm), ELS and MS (ESI pos and neg). HPLC sample preparation: 10 µl sample of each fraction was diluted 10x with $H_2O$/acetone 4:6 (90 µl). Injected 10 µl. For negative control, analysis of a sample obtained from the unspent media was performed in parallel. Fractions containing the target Syringomycin E were pooled and further separated on RP-C18 column (Phenomenex Kinetex XB-C18, 250 × 21.2 mm, 5 µm) using preparative HPLC (Agilent 1260). The analyte was eluted using gradient of B (MeCN + 2.0% FA) against A (MQ $H_2O$ + 2.0% FA). The method started at 30% B for 2 min, then gradient to 98% over 10 min, hold for 2 min, back to 30 % over 0.1 min and hold 4.9 min, all at flow rate 20 ml/min with detection by UV (210, 230, 250 and 400 nm), ELS and MS (ESI pos and neg). SIM trigerred fraction collection was employed monitoring m/z 1225.6 $[M+H]^+$. Fractions containing Syringomycin E were pooled and evaporated using Genevac EZ-2 Elite (HPLC lyo setting). Isolated yield after lyophilisation 11.7 mg.

Purity of the Syringomycin E sample (Fig. S3A) was determined to be 94% by HPLC (Dionex Ultimate 3000). The purity check was performed on a C18(2) column (Phenomenex, Luna 100 Å, 250 x 10 mm, 5 µm) using gradient of B (MeCN + 0.1% TFA) against A (MQ $H_2O$ + 0.1% TFA) at flow rate 4 ml/min. The method started at 30% B for 4 min, then gradient to 98% B over 22 min, hold for 4 min, then back to 30% B over 1 min and equilibrate for 4 min with UV detection at 210 and 265 nm. Samples were applied in water/acetone 4:6 (SRE c ~ 2 mg/ml). HR-MS (Fig. S3BC) on Synapt G2-Si for $C_{53}H_{86}ClN_{14}O_{17}$ $[M+H]^+$ = $C_{53}H_{86}ClN_{14}O_{17}$ calculated 1225.5980, observed 1225.5988, error 0.65 ppm.

**Bioactivity assay for purified SRE**

During SRE purification, fractions were tested for possible bioactivity against the filamentous eukaryotic microbe *Phytophthora infestans.* These assays were based on similar SRE purification/bioactivity experiments that used fungi rather than oomycetes. To test for bioactivity, we prepared potato dextrose agar (PDA; Formedium PDA0102) plates by roughly finding the centre of the plate and spotting appropriate buffer/fractions 3 cm away on each side from the centre. Plates were allowed to dry in a laminar flow hood, then an agar hyphal plug containing *P. infestans* (Pi88069) grown on PDA (7-day-old culture) was placed (hyphal side down) on the plate. Plates were sealed with micropore tape and grown at 18 $^oC$ in the dark for 6 days.
